# Dermal wound healing contribution of aqueous extracts of *Acalypha indica, Calotropis gigantea, Bacopa monnieri* and their combination

**DOI:** 10.1101/2024.01.18.576335

**Authors:** Habibu Tanimu, Ravindra Zirmire K, Colin Jamora, Parimala Karthik, O.S Bindhu

**Affiliations:** Department of Chemistry and Biochemistry, School of Sciences, Jain (Deemed to be University) Bangalore, Karnataka, India - 560027; IFOM-inStem Joint Research Laboratory, Centre for Inflammation and Tissue Homeostasis, Institute of Stem Cell Sciences and Regenerative Medicine, Gandhi Krishi Vigyana Kendra (GKVK) Campus, Bellary road, Bangalore, Karnataka India, −560065

**Keywords:** Angiogenesis, collagen remodelling, epidermal differentiation, polyherbal combination, re-epithelialisation, wound healing

## Abstract

Wound healing is a complex process that requires a well-orchestrated integration of an array of molecular events such as cell migration and proliferation, deposition and remodeling of extracellular matrix components for restoring the structural and functional integrity of the tissue injured. Ayurveda suggests wound healing herbs can achieve enhanced therapeutic effect with reduced toxicity when they are optimally combined in a specific ratio as polyherbal formulation (PHF). The present study was aimed to evaluate the combinatorial wound healing efficacy (*in vivo* wound closure and histological changes) of aqueous extracts of three medicinal plants (*Bacopa monnieri*, *Acalypha indica* and *Calotropis gigantea*). This study also explored how the combination influenced the overall quality of healed wound. Individual wound closure kinetic performance of aqueous plant extracts in C57B/6J mice was assessed using safe concentrations obtained from human adult dermal fibroblast viability assay. The aqueous plant extract combination optimized using response surface methodology was tested for *in vivo* wound closure effectiveness. Quality of healed wound was assessed via Hematoxylin & Eosin and immunohistochemical staining of markers (K1, K5, Loricrin, Ki67, CD31 and collagen1). The combination treatment *(B.monnieri*-15μg/ml, *A.indica*-11.59μg/ml, *C.gigantea*-1μg/ml) contributed to faster wound closure (11 days), improved collagen type I remodeling and angiogenesis, complete re-epithelialization, similar epidermal differentiation pattern as that of individual and control treatments. Ki67 staining revealed no significant increase in cell proliferation in combination compared to individual and control. Findings from the study validates the polyherbal combination’s impressive capability to promote wound healing.

## 1. INTRODUCTION

The complex process of wound healing with sequential but overlapping events necessitates the coordinated synchronization of different cell types and their products.^1^ Under homeostatic condition, the skin’s outer impermeable epidermis, middle dermis with rich extracellular matrix (ECM) and vasculature along with lower adipose tissue are engaged in maintaining its structural integrity.^2^ This protective effect is further complemented by constant surveillance from resident immune cells that are present in each layer of the skin.^3^ During injury, there develops a need for coordination between multiple cell types within these three skin layers at precise stages (hemostasis, inflammation, angiogenesis, growth, re-epithelialization, and remodeling) to bring about healing.^4^ Tissue regeneration and repair has been considered vital in this process that integrate several dynamic changes involving soluble mediators, blood cells, extracellular matrix production and parenchymal cell proliferation.

Though the wound repair program is remarkably efficient, there are instances such as aging, disease, and magnitude of the tissue damage that require intervention to facilitate and/or accelerate the healing process. Medicinal plants have been in use over years and still maintain an important role as first-line therapy for treating inflammation, burns, ulcers, and surgical wounds in different parts of the world.^5^ Diverse natural bioactive compounds present in them demonstrated their potential in accelerating the process of wound healing and regenerate tissue at the wound site. Research in this domain has furthered the relevance of phytotherapy and phytochemicals’ contribution on chemical signaling, organization of cells and extracellular matrix (ECM) modification in lesioned skin.^6^

The multifactorial nature of the wound healing program has given a pointer to why single agent treatment or use of individual plants for wound treatments is inadequate to produce the desired healing result. Use of medicinal plant combinations in the form of polyherbal formulation /combination appears to have a lot of promising potentials in wound treatment and management by decreasing closure time and/or increasing the quality of healed wounds over individual plant treatment.^7,8^ The availability of multitargeted groups of phytoconstituents has been considered as the cause for the improved efficacy possessed by these combinatorial preparations.^9,10^

Three important medicinal plants (*Bacopa monnieri*, *Acalypha indica*, *Calotropis gigantea*) have been selected to assess their combinatorial wound healing efficacy. The choice of these medicinal plants was made on the basis of their reported individual wound healing contribution at various phases of healing process. *Bacopa monnieri* has been reported to moderate the secretion of collagen type I and III, improve collagen fibre organization and remodeling.^11,12^ *Acalypa indica* was found to exhibit angiogenic activity by enhancing Vascular Endothelial Growth Factor (VEGF) and up-regulate selected pro-wound healing cytokines.^13–15^ Topical application of *Calotropis gigantea* on wound was reported to modulate type I collagen secretion and reorganization during wound healing.^16^

Considering the knowledge pertaining to these plants from the ethnopharmacological literature along with their crucial wound healing contribution evidences individually, this study tried to investigate their combinatorial efficacy towards *in vivo* wound closure kinetics and the key histological changes that promote effective healing of wounds. This study also explored how the combination influenced the overall quality of healed wound.

## 2. MATERIALS AND METHODS

### 2.1 Ethical Approval

Animal work conducted at the NCBS/inStem Animal Care and Resource Centre was approved by the inStem Institutional Animal Ethics Committee following the norms specified by the Committee for the Purpose of Control and Supervision of Experiments on Animals (Government of India) - approval number INS-IAE-2019/06 R1_ME). All experimental work was approved by the inStem Institutional Biosafety Committee (IBSC) and by research committee of Jain deemed to be University (USN:172/PHDBC06).

### 2.2 Plant collection, authentication and extract preparation

Fresh aerial plants of *B. monnieri* (Brahmi) and *A. indica* (Indian acalypha) were procured from Department of Horticulture, University of Agricultural Sciences (GKVK), Bangalore, Karnataka, India and were authenticated by Professor M. Vasundhara, Head, Department of Horticulture from the same institute (voucher numbers 65 and 50 respectively). *C. gigantea* (Giant shallow wort) was collected from outskirts of South Bangalore and was authenticated by Dr. Shiddamallayya N. Assistance Research Officer (Botany), National Ayurveda Dietetics Research Institute (NADRI), Jayanagar, Bangalore, Karnataka, India (voucher number SMPU/NADRI/BNG/2010-11/490).^17,18^ The plants were thoroughly washed, shade dried, and made into a coarse powder using an electric homogenizer. Aqueous extract was made by dissolving 25 g of the powdered plant in conical flask containing 250 ml of distilled water. The extracts were filtered after keeping on a shaker for 24 h at room temperature using Whatman No. 1 filter paper. The filtrates were lyophilized using Labconco Freeze dryer by LABQUIP India Private limited (Hyderabad, India). The dried extracts were packed in a closed container and stored at −20°C for further analysis.^19–21,17^

### 2.3 Chemicals and reagents

All chemicals and reagents used in this research were of analytical grade and were procured from HiMedia laboratories and Thermo Fisher Scientific India Pvt Ltd Mumbai, India.

### 2.4 Cell culture and *in vitro* cell viability assay

Human adult dermal fibroblasts (HADF) (Lonza CC-2511) were cultured in Dulbecco’s Modified Eagle Medium (DMEM) with 10% Fetal Bovine Serum (FBS), antibiotic (Pen-Strep), Non-Essential Amino Acid (NEAA) and sodium pyruvate. The cells were kept in an incubator set at 37°C with humidified atmosphere of 95% air supplied with 5% carbon dioxide (CO_2_).^22^

Plant extracts were tested for their cell viability using 2-(4-iodophenyl)-3-(4-nitrophenyl)-5-(2, 4-disulfophenyl)-2H-tetrazolium, monosodium salt (WST-1 assay) (Roche, Manheim, Germany, Cat. #5015944001) according to the manufacturer’s instructions. Briefly, HADF cells were seeded in 96 well plates at a density of 2000 cells/well in 10% serum medium and incubated for 24 h. Different concentrations of plant extracts were added to the cells with fresh medium supplemented with 1% FBS and further incubated for 72 h. The media was aspirated out and 10% WST-1 in 1% serum medium was added to each well and incubated again for 2 h to allow colour development. Absorbance was measured using microplate reader (Varioskan^TM^ lux multimode plate reader, Thermo Scientific United states) with test and reference wavelengths at 450 nm and 650 nm respectively.^23,24,22^

### 2.5 Wound closure kinetics of individual plant extracts

Plants extracts were individually tested for wound closure kinetics based on the optimized concentration from *in vitro* cell viability assay results as described by^25^ Biswas et al., (2017) and ^26^Gund et al., (2021). Healthy 8 weeks old male Black 6 Jackson mice (C57B6/J) weighing 22-25 g were used. Animals were divided into four groups of three each (group I - vehicle control, group II – *B.monnieri*, group III - *A.indica*, group IV - *C.gigantea*).^27^ The mice were anesthetized with isoflurane gas after which full thickness excision wounds were created on the dorsal side. The optimized concentrations of the plant extracts were applied topically once daily until wound closes. Distilled water treated group served as control.

The excision wounds’ contracting diameter (mm) were observed to macroscopically evaluate the wound healing features and kinetics. Diameter of the wounds was photographed using digital camera (Nikon D70, Tokyo, Japan) daily from day one till the wound closes. Measurement and analysis were done using ImageJ software version 1.53 (Maryland, USA). Percentage wound closure was calculated using the following formula:

Percentage wound closure (%) = A Day 0 – A Day X/ A Day 0 *100

Where, Day X = Day of evaluation and A = diameter of the wound

### 2.6 Wound closure kinetics of the plant extract combination

The aqueous plant extract combination optimized using response surface methodology (RSM) (central composite design using a 20 factorial and star design with three central points, Design Expert 12 software - trial version) was tested for *in vivo* wound closure kinetics.^25,26^ C57B/6J mice were anesthetized with isoflurane gas post which full thickness excision wounds were created on the dorsal side of each mouse following the same procedure explained previously under section 2.5.

### 2.7 Tissue processing and embedding

After the wound was closed, the mice were immediately euthanized with 100% CO_2_ inhalation. The granulation tissues (1cm×1cm) were removed using a sterile biopsy punch, and fixed in Optimum Cutting Temperature (OCT) solution (Leica, catalogue number: 14020108926). The fixed tissue was sectioned (12 µm thick) and stained appropriately for microscopic evaluations.^26^

#### 2.7.1 Hematoxylin and Eosin (H&E) staining

Histological analysis of wound skin sections was performed by H & E staining. Skin sections stored at −80^°^C were thawed and fixed with 4% formaldehyde for 10 min. The slides were washed three times (5 min each) with phosphate buffered saline (PBS) and was given 10 dips in hematoxylin solution followed by 10 dips in tap water to remove excess hematoxylin from the sections. The slides were then immersed in eosin solution for a few seconds followed by a dip in distilled water. After draining the water, a few drops of 80% glycerol were applied on the sections followed by mounting with a coverslip.^28,29^

#### 2.7.2 Immunohistochemical (IHC) analysis

OCT embedded wounded skin sections were processed and immunohistochemistry was performed as described by^1^ Lee et al., (2017) and ^30^Du et al., (2010). Quantification of Ki67 was done according to^31^Arwert et al., (2010). Slides were thawed in a humid chamber for 10 min. Tissues were fixed with 4% Paraformaldehyde (PFA) for 10 min after which they were washed with 0.1% Phosphate buffered saline with Tween-20 detergent (PBST) two times for 5 min each. PAP pen was used to make a mark around the tissue on the slide to prevent spilling of blocking solution and antibodies. Blocking solution was added, incubated at room temperature for 1 h and removed. Unlabeled specific primary antibodies dissolved in blocking solution were added and incubated at 4°C overnight. The primary antibody was washed with 0.1% PBST three times 5 min each. Fluorescent labelled secondary antibodies were added and incubated for 20-25 min at room temperature and removed by washing with 0.1%PBST (three times for 5 min each). For nuclear staining, 4’,6-Diamidino-2-Phenylindole (DAPI) or Hoechst were used. The sections were mounted with anti-fading solution, Mowiol. Images of the stained sections were taken immediately using inverted microscope (Olympus DP80, IX73). Freely available ImageJ software (version 1.8.1, Public Domain, Madison, WI, USA) was used for image analysis.^32^ All the antibodies and their corresponding dilutions used in this study are presented in supplementary material-Table 1.

#### 2.7.3 Statistical analysis

All the experimental data of replicates from three independent experiments were reported as mean ± standard error of mean (SEM). Statistical significance in the difference between various treatments and respective controls was evaluated using two-way ANOVA. Tukey’s post hoc test was used for multiple comparions. Graphpad Prism (Version 8.0) was used and P value ≤ 0.05 was considered statistically significant.

## 3. RESULTS

### 3.1 *In vitro* cell viability and *in vivo* wound closure concentration of *B.monnieri, A.indica and C.gigantea*

*In vitro* tolerable concentration of the plant extracts was determined on primary human dermal fibroblasts (HDF) using WST-1 assay. HDF cells were exposed to increasing concentrations of the plant extracts for 72 h and their effect on cell viability was evaluated. *B.monnieri*, *A.indica* and *C.gigantea* presented 5-20 µg/ml, 10-80 µg/ml, and 0.2-1 µg/ml as tolerable range of viable concentrations respectively. The concentration of extracts used for treatment and their corresponding cell viability are presented in Figure 1A.

**FIGURE 1.**
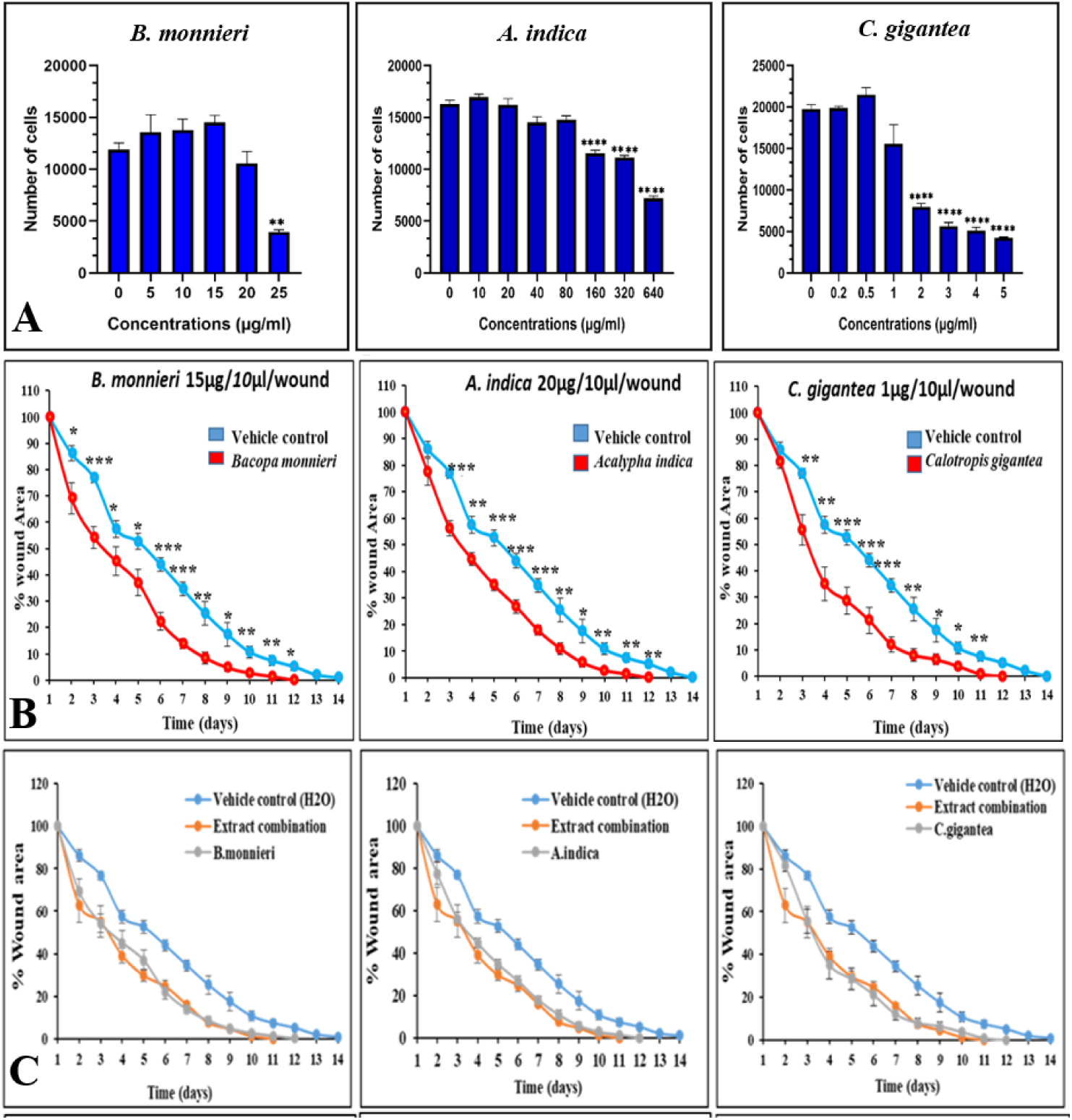
*In vitro* and *in vivo* effects of *B.monnieri, A.indica* and *C.gigantea* aqueous extracts. Effect of plant extract concentrations on the viability of HDF (A) Wound closure kinetics of plant extracts (B) Wound closure kinetics of plant extracts and their combinations (C). Vehicle control= Distilled water. Data presented as mean ± SEM of triplicates from three independent biological replicates. P values were calculated using two-way ANOVA. Tukey’s post hoc test was adopted for multiple comparison (ns = not significant, *P < 0.05, **P < 0.01, ***P<0.001, ****P<0.0001)

To further understand the *in vivo* wound healing efficacy of these concentrations, wound closure kinetic assay was set up on C57B6/J wild type mice. Concentrations that exhibited the fastest wound closure were noted (Figure 1B). All three plant extracts *(B.monnieri*, *A.indica* and *C.gigantea* at 15 µg/ml, 20 µg/ml and 1 µg/ml respectively) exhibited improved wound closure contribution (complete closure 12 days post wounding [dpw]) compared to the vehicle control treated group (complete closure 14 dpw).

### 3.2 Optimization of plant extract concentrations to evaluate their combinatorial efficacy on *in vivo* wound closure

Central composite design (CCD) of Response Surface Methodology (RSM) was adopted to optimize the required concentration of the plant extracts towards the Polyherbal Combination (PHC) preparation to further understand their combinatorial efficacy on *in vivo* wound closure kinetics. This statistical tool was used to study the nature of interactive effect of respective plant extracts’ concentration (independent variable) and their effect on wound closure time in days (dependent variable). The range of independent variables was selected based on preliminary *in vitro* cell viability results (Supplementary file-Table 2). From the twenty independent runs carried out, the optimum concentrations of individual plant extract in combination for improved wound closure was found to be: *B.monnieri* (15 µg/ml), *A.indica* (11.591 µg/ml) and *C.gigantea* (1 µg/ml) (Supplementary file-Table 3). Adequacies and fitness of the chosen quadratic model was verified by ANOVA results (Supplementary file-Table 4) where the model appeared significant with a non-significant lack of fit. The relationship between the independent variables was also obtained from the regression analysis result (Supplementary file-Table 5). The relationship of the three variables *B.monnieri, A.indica and C.gigantea* for wound closure rate in terms of coded values is given by the following equation:

Y = 12.47+ (0.0732*A) + (0.0882*B) – (0.2696*C) + (0.1250*AB) + (0.2500*BC) + (0.7785*A^2^) + (0.2482*C^2^). Where Y in the equation represents the quadratic model of the response (wound closure rate), the notation A, B, C, AB, BC, A^2^ and C^2^ represent the input factor values of *B.monnieri, A.indica* and *C.gigantea*.

Results of *in vivo* wound closure kinetics of this extract combination [*B.monnieri* (15 µg/ml), *A.indica* (11.591 µg/ml) and *C.gigantea* (1 µg/ml)] exhibited faster wound closure rate than individual plant extracts (Figure 1C). The extract combination-treated group closed on day 11, while each of the individual plant-extract treated group closed on 12 dpw and the vehicle control-treated wounds closed 14 dpw. Representative images of wounds created on the mice from day 1 till the day of wound closure is presented in Figure 2.

**FIGURE 2.**
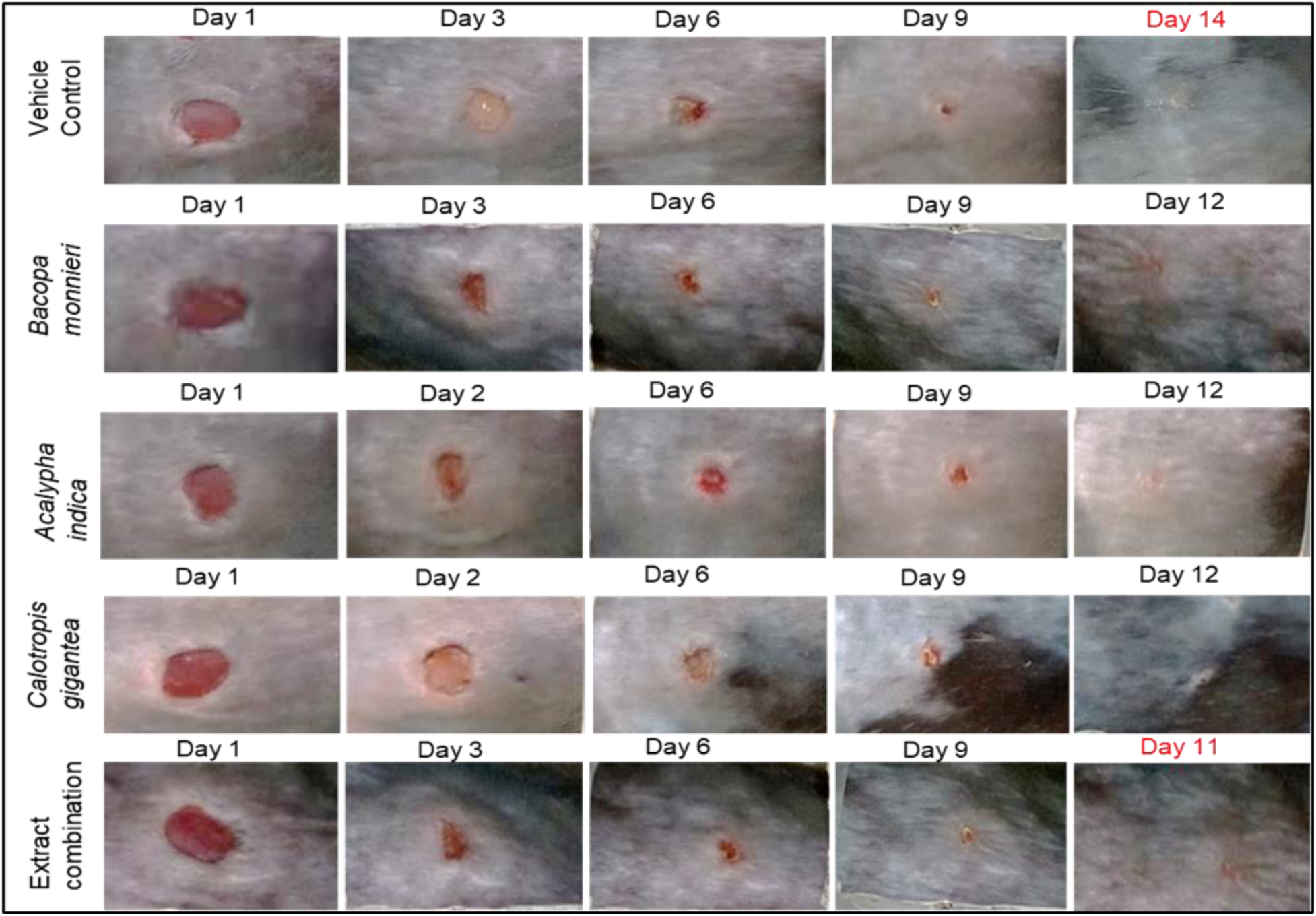
Healing progress of wounds after application of the plant extracts and their combination on wild type C57B6/J mice. Day 1 indicates the wounding day from where wound closure rate was measured daily till the wound closes.

### 3.3 PHC treatment improved histological state and architecture of the wounds

To assess whether the quality of wound repair was compromised for the sake of faster closure, histological examination of the wound tissue was conducted. Mice skin sections were harvested immediately upon complete wound closure (14, 12 and 11 dpw for control, individual extract and combination treatments respectively) and processed for H & E staining to evaluate the architecture and state of the wound repair process following treatment with individual plant extracts and their combination compared to vehicle control. The histology of all wounds (plant extract treated groups and the control) were examined (Figure 3A-E). An increase in both epidermal and dermal thickness is indicative of a pathological wound healing response. Quantification data revealed higher dermal thickness by the *B.monnieri* treated group (Figure 3F) and higher epidermal thickness by *A.indica* treated group (Figure 3G) compared to other treatments in the study. The interesting result is that these abnormalities are abolished in the combination.

**FIGURE 3.**
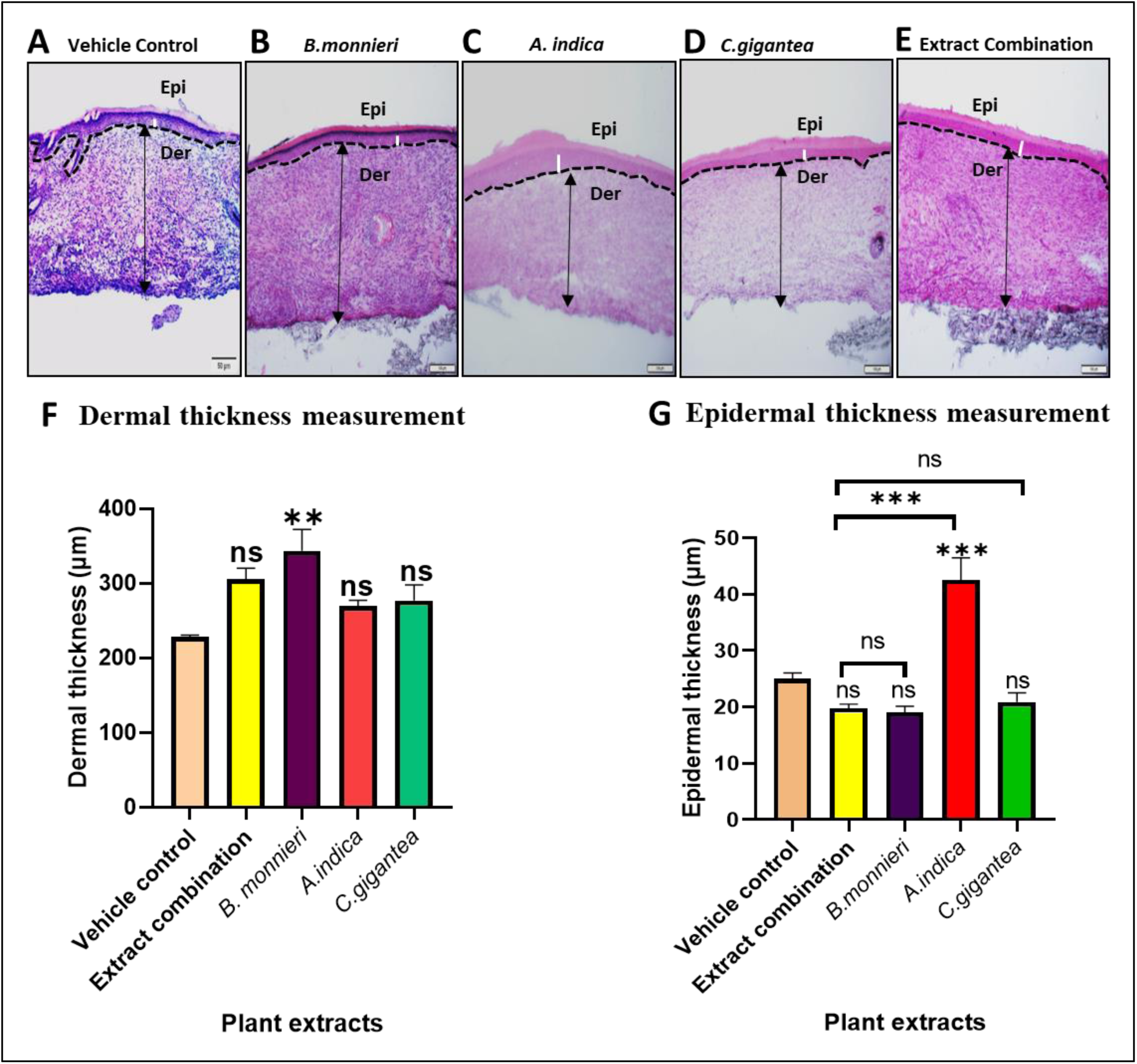
Photomicrographs of H&E-staining of histological sections representative of wound healing area in mice from different groups and the corresponding epidermal and dermal thickness measurement. White line represents epidermal thickness while black arrow denotes dermal thickness. Black dotted line indicates the epidermal–dermal interface. Staining was carried out on OCT embedded tissue sections. Data presented as mean ± SEM of triplicates from three independent biological replicates. P values were calculated using one way ANOVA. (ns= not significant, **\*\*P < 0.01, ***P<0.001)**, Tukey’s post hoc test was adopted for multiple comparison. Epi=Epidermis, Der=Dermis. Scale bar-50µm.

### 3.4 Extract combination did not affect epidermal layer’s differentiation

To understand whether epidermal differentiation was affected by the treatment with extract, a molecular expression profile of the healed tissues by immunohistochemical staining of specific epidermal markers was carried out. The markers such as anti K5 antibody (staining basal keratinocytes), anti K1 antibody (staining terminally differentiated keratinocytes) and anti Loricrin antibody (staining Loricrin, a major protective protein component produced by terminal keratinocytes) were used to assess the epidermal differentiation (Figure 4). Individual plant extract and their combination treated groups exhibited a similar epidermal differentiation pattern as the control with orderly restoration of all epidermal layers of the re-epithelialized wounded tissue.

**FIGURE 4.**
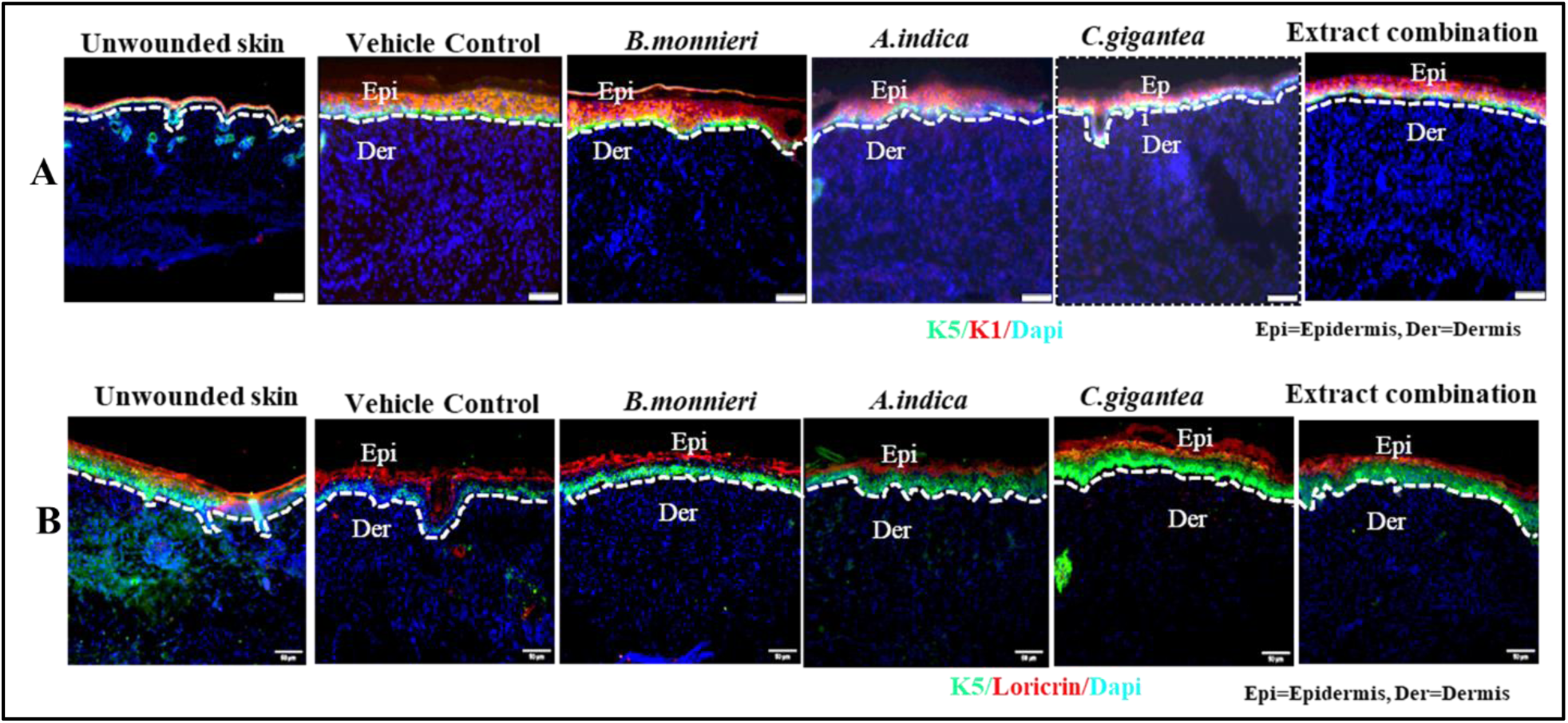
Expression of epidermal markers. (A) Immunostaining of unwounded, vehicle control, individual extracts and extract combination with anti K5, K1 and Dapi. (B) Immunostaining of unwounded, vehicle control, individual extracts and extract combination anti K5, Loricrin and Dapi antibodies. The white dotted line represents the epidermal–dermal interface. Staining was carried out on OCT embedded tissue sections. Epi=Epidermis, Der=Dermis. Scale bar-50µm.

### 3.5 Extract combination did not significantly increase cell proliferation

Proliferation is one of the most important phases of wound healing. For quality performance of wound healing, proliferation has to be maintained in a regulated manner. Excessively low and high cell proliferation during wound healing can lead to conditions such as delay in wound healing and scarring respectively. Wound tissue sections stained with marker of proliferation, Ki67, showed that extract combination did not significantly increase cell proliferation compared to individual plant and vehicle control treated groups (Figure 5).

**FIGURE 5.**
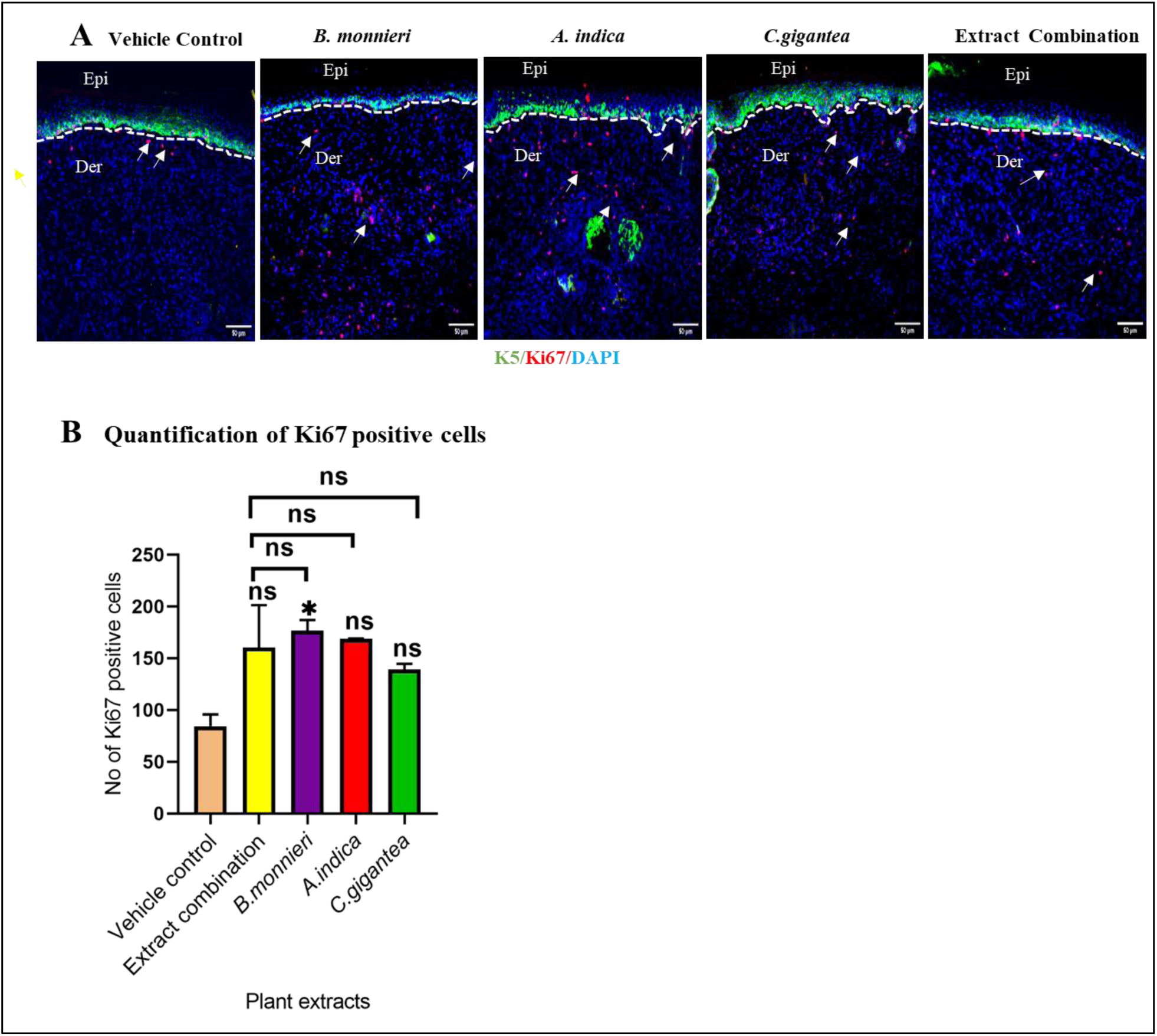
Immunohistochemical staining for marker of cell proliferation. (A) Skin wound sections treated with plant extracts showing Ki67 positive cells. (B) Quantification of ki67 positive cells. White dotted line represents the epidermal–dermal interface. White arrows indicate Ki67 positive cells. All staining is done on OCT embedded tissue sections. Data presented as mean ± SEM of triplicates from three independent biological replicates. P values were calculated using one way ANOVA. [ns (above the bars) =not significant when compared to the control group, ns (above the brackets) =not significant performance of individual extract compared to that of combination. *P < 0.05). Tukey’s post hoc test was used for multiple comparison. Epi=Epidermis, Der=Dermis. Scale bar-50µm.

### 3.6 Extract combination significantly increased angiogenesis

An important factor determining the functionality of repaired skin is restoration of proper vascularization of the tissue. Restoring the vascular system of the skin is necessary because newly formed tissues need to get supply of nutrients. Staining of the wounded skin sections with endothelial cell marker have shown that extract combination treated group exhibited significant vasculature (P<0.01) compared to individual extracts and vehicle control (Figure 6).

**FIGURE 6.**
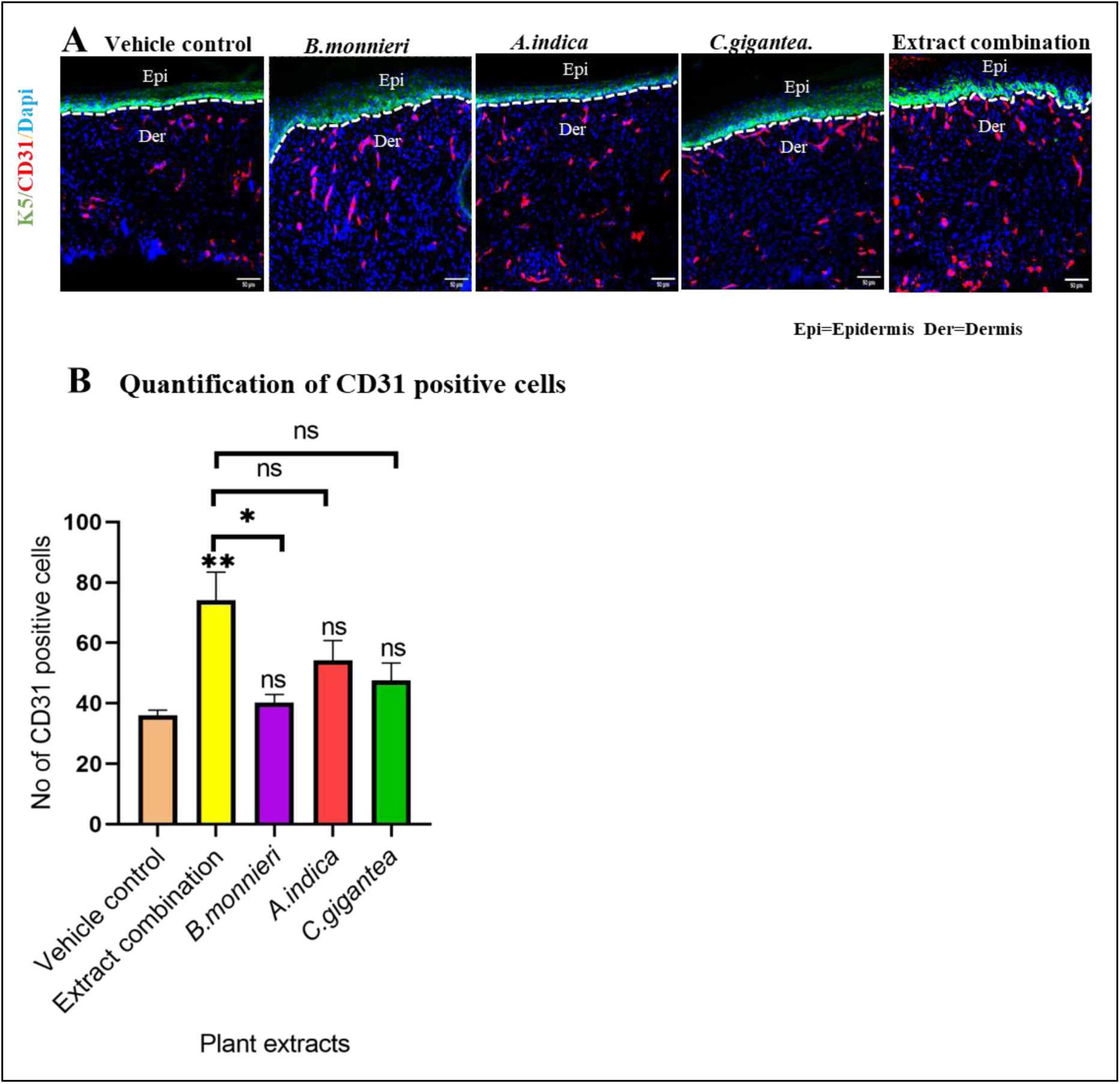
Immunohistochemical staining of endothelial cells using CD31 antibody. Compared to the control group, sections from individual plant extract and combination treated tissues exhibited higher number of endothelial cell formation with combination displaying a significant difference (P<0.01). (A) Tissue sections from different study groups immunostained with anti CD31 antibody and (B) corresponding quantification of CD31 positive cells. Data presented as mean ± SEM of triplicates from three independent biological replicates. P values were calculated using one way ANOVA. [ns (above the bars) =not significant when compared to the control group, ns (above the brackets) =not significant performance of individual extract compared to that of combination. *P < 0.05, **P<0.01). Tukey’s post hoc test was adopted for multiple comparison. White dotted line represents the epidermal–dermal interface. Epi=Epidermis, Der=Dermis. Scale bar-50µm. All staining is done on OCT embedded tissue sections.

### 3.7 Extract combination improved collagen type I remodeling

A major component determining the structural integrity of skin is the proper formation of extracellular matrix in the dermis, of which collagen is the predominant component. To check for collagen I expression, immunohistochemical staining with anti-collagen I antibody was performed. Findings indicated that extract combination exhibited more of the fibril organization of collagen I that is found in the unwounded skin compared with the individual plant extract and vehicle control treated groups (Figure 7).

**FIGURE 7.**
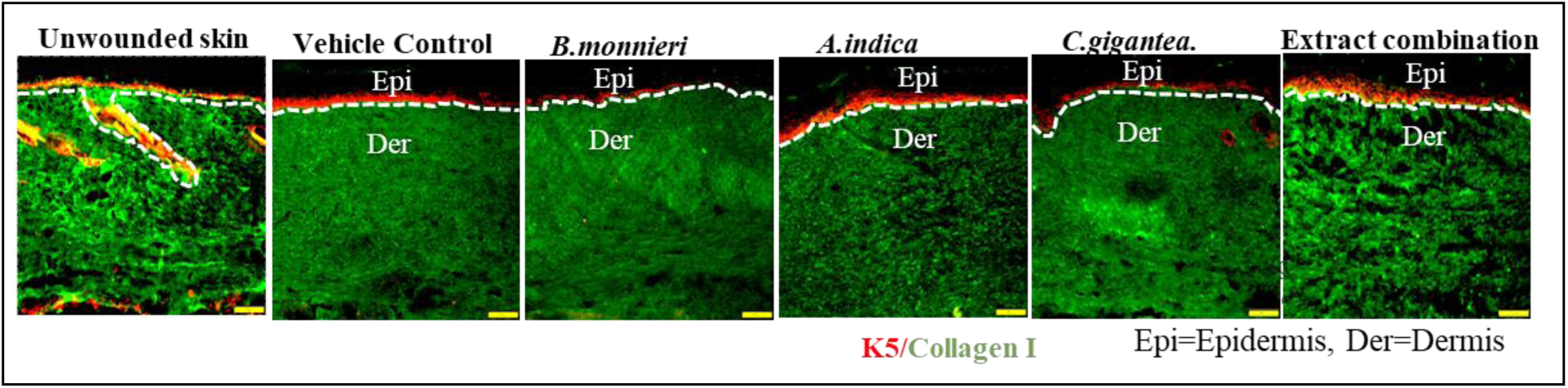
Immunohistochemical staining of collagen I. (A) Unwounded skin tissue section. (B) Individual plant extracts and combination treated wound tissue sections. Extract combination showed improved collagen type I fibril organisation similar to unwounded skin section. White dotted line represents the epidermal–dermal interface. All staining was done on OCT embedded sections. Epi=Epidermis, Der=Dermis. Scale bar - 50µm.

## 4. DISCUSSION

Chronicity of wound is an enduring challenge for patients and health care professionals alike globally. Failure of an orderly progression cascade of wound healing or stalling in the inflammatory phase are considered as the primary reasons that prevent wound from progressing further into the healing continuum. This will invariably result in long-term impaired healing process, in which proper anatomical and functional results are not achieved within a designated period. Plant extracts and their combination have been demonstrated to accelerate wound healing via multiple pharmacological targets.^33^. They improve the wound condition and prevent chronicity by interfering with some pathophysiological effects (reduction of inflammatory cell influx, promotion of new blood vessel formation, favoring fibroblastic proliferation and reepithelization) either independently or in combination. Polyherbal approach advocates the use of multiple herbs rather than single herbal formulation. This system utilizes the concept of positive herb-herb interaction where multiple herbs in a single formulation help in striking many potential targets at the same time leading to improved outcome. The wound healing aspect happens to be the most recognized application of polyherbal formulations amongst its other reported health benefits.^34^ Ayurvedic and Indian traditional medicine consider *B. monnieri, A. indica* and *C gigantea* as highly regarded medicinal plants owing to their broad-spectrum bioactivities. Existing literature reveals that extracts or active constituents isolated from these plants have wound healing potential. *B.monnieri* and *C.gigantea* latex were reported to exhibit faster wound closure rate as compared to nitrofurazone (0.2w/v), a standard drug employed for wound healing.^11,35^ *A.indica* was reported to increase the rate of wound contraction and faster re-epithelialization compared to betadine ointment used as positive control.^36^ *Bacopa monnieri, Acalypha indica* and *Calotropis gigantea* were selected for investigating their combinatorial efficacy on the architectural changes in the histology of wound tissue post healing in comparison with that of the individual extract treatment groups. Changes with respect to skin tissue histology, epidermal differentiation, epidermal & dermal thickness, cell proliferation tendency, angiogenesis and collagen type I remodeling were assessed using suitable assays. From the cell viability results (Figure 1A), the following concentrations were considered for assessing *in vivo* wound closure contribution by *Bacopa monnieri* (15 µg/ml), *Acalypha indica* (20 µg/ml) and *Calotropis gigantea* (1 µg/ml). Mouse skin samples were processed and histological analysis by H&E staining was carried out to assess the state of wound repair process after healing following treatment with individual plant extracts and their combination compared to vehicle control treated group.^30^ Complete wound closure was found to be 11 dpw, 12 dpw, 14 dpw for combination, individual extract and vehicle control treatments respectively. The wounded skin tissues stained with H & E (Figure 3) revealed that there is complete re-epithelialization of wounds for all the treated groups. Epidermal and dermal thickness were quantified to assess indication of pathological condition, if any, that might have ocurred during the wound healing process.^30^ Increase in epidermal thickness is indicative of hyperproliferation and activation of keratinocytes that may contribute to pathological condition interfering with the healing of wounds.^37^ Excessive increase in epidermal thickness is a pathlogical condition that occurs when normal wound healing process gets pertubed. This situation appears due to disproportionate production and accumulation of ECM during woud healing process leading to scarring, a pathological condition that is characterised by tissue hardening and limited function.^38^ All the individual plant extracts and their combination showed complete re-epithelialization. Our finding in this experiment indicates that extract combination closes the wound faster than individual plant extracts and the vehicle control. To understand whether quality is compromised to speed, epidermal thickness (Figure 3G) as well as dermal thickness (Figure 3F) were quantified. Despite the fact that extract combination significantly enhanced the wound closure rate compared to their individual counterpart and vehicle control, epidermal thickness did not seem to significantly increase. The extract combination appears to decrease the wound closure time through a synergistic or complementary interaction rather than additive as *A.indica* has shown a significant (P<0.001) increase in epidermal thickness. Similar trend was observed where the dermal thickness of extract combination treated group did not show a significant increase despite the significant increase (P<0.01) in that of *B.monnieri* treated group.

For wound to be considered properly healed, there must be proper epidermal layer differentiation as much as possible to the level of unwounded skin after wound healing.^39^ Markers that give information on structural intactness of the epidermis such as K5 which stains the basal epidermal layer, K1 which stains the terminal keratinocyte layer and loricrin which is a structural protein produced by terminal keratinocytes are studied.^1^ All the plant extracts individually and in combination did not seem to affect the epidermal differentiation as all the above markers’ staining details suggest the epidermal layers are back to normal as compared to unwounded skin section and vehicle control treated wound. This finding suggests that the faster wound closure activity of extract combination has not compromised the quality of the epidermal layer differentiation. Epidermal layer restoration observed in this study is in line with observation reported by^40^, where a PHF containing *Piper betle*, *Curcuma longa*, *Aloe vera* and *Thymus vulgaris* enhanced wound closure and restored back intact epidermal layers in an *in vivo* experiment with wistar albino rat. Regenerative and wound repair process is a complex phenomenon involving numerous cell types. These cells carry out coordinated interaction to restore the lost tissue during wound healing process. Well-organized wound repair process depends largely on the synchronized interactions between hemostasis, inflammatory, proliferative and remodeling phases of the wound-healing response.^4^ After hemostasis which last for just a few seconds, inflammatory and proliferative phase immediately begin where those large number of cells increase in number to facilitate the wound repair process. Despite the importance of the proliferative phase of wound healing, it has to be maintained in a homeostatic level as excessively low proliferation rate will lead to delay in wound healing while hyper proliferation of cells can lead to pathological condition such as chronic wound or cancer.^41^ Wound healing experiment carried out in this study indicated that extract combination did not significantly increase cell proliferation compared to the vehicle control. Although all the plant extract treated groups showed an increase in cell proliferation compared to the vehicle control, only *B.monnieri* appeared to have exhibited significant (P<0.05) cell proliferation. This finding suggests that the faster wound closure activity shown by the extract combination could be through other molecular mechanism not as a result of increase in cell proliferation during the wound healing process. This result is in line with the findings of a previous work where *in vitro* antiproliferative activity of a PHF was reported for the treatment of cancer.^42^ During wound healing, restoring vascular system of the skin is necessary because newly formed tissues need to get supply of nutrients to be functional for improved wound healing. Several growth factors such as Vascular Endothelial Growth Factor (VEGF), Platelet Derived Growth Factors (PDGF) and Fibroblast Growth Factor-basic (bFGF) have been reported to be involved in initiating the neovascularization process during wound healing.^43^ Extract combination treated group exhibited significant vasculature compared to the individual extract treated group and vehicle control (Figure 6). This finding suggests that the faster wound closure displayed by extract combination treated group could be an outcome of the increased neovasculization/angiogenesis. This result is in line with findings of ^44^, where a PHF containing extracts of *Vitex negundo* L., *Emblica officinalis* Gaertn, and *Tridax procumbens* L. was reported to significantly increase the density of blood vessels compared to the control using Chorioallantoic Membrane assay.

Collagen I, the most abundant collagen in all vertebrates, plays a critical role in giving mechanical strength to the tissue during and after wound healing.^45^ It is one of the major components of ECM, produced and deposited by fibroblasts during wound healing, and its role in re-epithelialization along with tissue homeostasis has also been reported.^40,46^ The sequential appearance and remodeling of ECM, particularly collagen type I and III, play a decisive role in contributing to the strength of a healing wound. Type I collagen deposition and remodeling was recognized as a contributor in increasing tensile strength of the wound.^47^ Many natural extracts explored towards wound healing also induced the expression and improved morphology of collagen type I in the healing wound tissue.^48^ Inadequate ECM production leads to pathological condition that results in premature aging.^49^ Excessive ECM production brings about several pathologies such as scarring of different types, scleroderma, fibrosis and organ failure. Findings from this study has indicated that extract combination has improved collagen I remodeling effect compared with that of the individual plant extract treated group and vehicle control (Figure 7). Similar trend was observed when a PHF containing *Piper betle*, *Curcuma longa, Aloe vera* and *Thymus vulgaris* was reported to moderate the synthesis and degradation of collagen I by increasing and proportionately decreasing its production at the beginning and end of wound healing process respectively.^40^ Improved wound healing properties of a polyherbal ointment via moderation of collagen secretion and anti-inflammatory activity was reported by^50^ Elzayat et al., (2018). Yet another study reported the accelerated wound healing activity of a PHF containing *Martynia annua* L. and *Tephrosia purpurea* via improved collagen remodeling and established molecular cross linking of collagen fibres.^51^ Results from this study pointed out the advantage of herbal combination in regulating the process of wound healing relative to the use of individual plant through various molecular mechanisms.

## ACKNOWLEDGMENT

The authors thank Dr. Leela Iyengar, Former adjunct Professor, Jain (Deemed-to-be University), Bengaluru, India and Chief Scientific Officer (Retd), I.I.T. Kanpur, India, for her suggestions and critical revision of the manuscript. Financial support was provided by Jain (Deemed-to-be University) through seed money project grant.

