## Supplementary material for "Dermal wound healing contribution of aqueous extracts of *Acalypha indica, Calotropis gigantea, Bacopa monnieri* and their combination"

**Table 1.** **Key resources used for histological and IHC analysis**

| **Designation of Animal/Antibodies Used** | **Sources or reference** | **Identifiers** | **Additional Information and Dilutions** |
| --- | --- | --- | --- |
| C57B6/JNcbs (mice) | Jackson Laboratory | Stock No. 000664 | Animals were originally purchased from Jackson Laboratory (Stock No. 000664) but were bred for > 5 generations in the NCBS animal facility |
| **Primary Antibodies** | | | |
| Anti-Ki67 (Rabbit) (Proliferation marker) | Abcam | ab16667 | 1:100 |
| Anti-Keratin 5 (Chicken) (Basal keratinocytes marker) | Jamora Lab generated |  | 1:200 |
| Anti-Loricrin (Rabbit) (Terminal keratinocytes marker) | Jamora Lab generated |  | 1:500 |
| Anti-CD31 (Rat) (Endothelial cells marker) | Abcam | AB28364 | 1:200 |
| Anti-Collagen 1 (Rabbit) | Abcam | AB21286 | 1:200 |
| DAPI (Nuclear stain) | Molecular Probes | AB228549 | 1:500 |
| **Secondary antibodies** | | | |
| Goat anti-Chicken Alexa fluor 488 | Invitrogen | A21202 | 1:200 |
| Donkey anti-Rabbit Alexa fluor 568 | Invitrogen | A10042 | 1:200 |
| Donkey anti-Rat Alexa fluor 488 | Invitrogen | A21208 | 1:200 |
| Goat anti-Chicken Alexa fluor 568 | Invitrogen | A11041 | 1:200 |
| Goat anti-Chicken Alexa fluor 488 | Invitrogen | A11034 | 1:200 |
| Donkey anti-Rabbit Alexa fluor 488 | Invitrogen | A21206 | 1:200 |
| Goat anti-Rabbit Alexa fluor 647 | Invitrogen | A21244 | 1:400 |

**Note:** Antibodies and other materials used for histological and IHC analysis. Dilution buffer is the blocking solution (Tris-Cl (pH 7.4) 10mM, MgCl_2_ 100mM, Tween 20 0.5% (v/v), Bovine serum albumin (BSA) 1% (w/v), Fetal bovine serum (FBS) 5% (v/v) (Sinha et al 2022)

**Table 2 Experimental range of concentrations for *B. monnieri, A. indica* and *C. gigantea* entered as coded values for optimization**

|  |  | Coded values | | |
| --- | --- | --- | --- | --- |
| **Plant extracts** | **Unit** | **-1** | **0** | **+1** |
| *Bacopa monnieri* | µg/ml | 10 | 15 | 20 |
| *Acalypha indica* | µg/ml | 15 | 20 | 25 |
| *Calotropis gigantea* | µg/ml | 0.5 | 1 | 1.5 |

**Note:** These concentrations are fed into design expert software as coded values where ‘0’ indicate actual concentration chosen and -1, and +1 indicate minimum and maximum concentration ranges.

**Table 3 Values of factors and response for optimization of the concentrations of *B*. *monnieri, A*. *indica and C*. *gigantea* plant extracts in CCD**

| **Run Number** | ***B*. *monnieri* µg/ml)** | ***A*. *indica* (µg/ml)** | ***C*. *gigantea* (µg/ml)** | **Wound Closure Time (days) Experimental** | **Wound Closure Time (days)**  **Predicted** |
| --- | --- | --- | --- | --- | --- |
| 1 | 20 | 15 | 0.5 | 14.00 | 13.45 |
| 2 | 15 | 20 | 1 | 12.50 | 12.57 |
| 3 | 10 | 15 | 1.5 | 13.50 | 12.91 |
| 4 | 15 | 28.409 | 1 | 13.50 | 12.74 |
| 5 | 15 | 20 | 1 | 13.50 | 12.57 |
| 6 | **15** | **11.591** | **1** | **11.00** | **12.20** |
| 7 | 15 | 20 | 1 | 12.50 | 12.57 |
| 8 | 15 | 20 | 0.159104 | 13.50 | 13.55 |
| 9 | 6.59104 | 20 | 1 | 14.00 | 14.09 |
| 10 | 15 | 20 | 1 | 12.00 | 12.57 |
| 11 | 10 | 25 | 1.5 | 13.00 | 13.24 |
| 12 | 15 | 20 | 1 | 12.50 | 12.57 |
| 13 | 20 | 15 | 1.5 | 13.50 | 12.81 |
| 14 | 15 | 20 | 1 | 12.50 | 12.57 |
| 15 | 23.409 | 20 | 1 | 14.00 | 14.34 |
| 16 | 10 | 25 | 0.5 | 13.00 | 13.38 |
| 17 | 20 | 25 | 0.5 | 13.50 | 13.78 |
| 18 | 20 | 25 | 1.5 | 13.50 | 13.63 |
| 19 | 10 | 15 | 0.5 | 14.00 | 13.56 |
| 20 | 15 | 20 | 1.8409 | 12.50 | 12.89 |

**NOTE:** Experimental and predicted values of interactions between the independent variables using 20 runs. Run number 6 showing the shortest wound closure time (11th day) was selected for *in vivo* combinatorial experiment.

**Table 4 Output of ANOVA using Quadratic Model of Response Surface for the plant extract combination**

| Source | **Sum of Squares** | **df** | **Mean Square** | **F-value** | **p-value** |  |
| --- | --- | --- | --- | --- | --- | --- |
| Model | 11.08 | 7 | 1.58 | 3.14 | 0.0396 | significant |
| A-*Bacopa monnieri* | 0.0732 | 1 | 0.0732 | 0.1451 | 0.7099 |  |
| B-*Acalypha indica* | 0.1062 | 1 | 0.1062 | 0.2105 | 0.6545 |  |
| C-*Calotropis Gigantea* | 0.9926 | 1 | 0.9926 | 1.97 | 0.1861 |  |
| AB | 0.1250 | 1 | 0.1250 | 0.2477 | 0.6277 |  |
| BC | 0.5000 | 1 | 0.5000 | 0.9909 | 0.3392 |  |
| A² | 8.82 | 1 | 8.82 | 17.48 | 0.0013 |  |
| C² | 0.8965 | 1 | 0.8965 | 1.78 | 0.2073 |  |
| Residual | 6.05 | 12 | 0.5046 |  |  |  |
| Lack of Fit | 4.55 | 7 | 0.6507 | 2.17 | 0.2056 | not significant |
| Pure Error | 1.50 | 5 | 0.3000 |  |  |  |
| Cor Total | 17.14 | 19 |  |  |  |  |

**Note:** The F-value of 3.14 and P-value of 0.0396 indicate that terms of the model are significant. The **lack of fit value** of 2.17 implies that it is not significant relative to the pure error. There is 20.56% chance that the lack of fit F-value this large could occur due to noise. Non-significant lack of fit is good.

**Table 5 Regression equation results of the plant extracts in terms of coded factors**

| **Factor** | **Coefficient Estimate** | **df** | **Standard Error** | **95% CI Low** | **95% CI High** | **VIF** |
| --- | --- | --- | --- | --- | --- | --- |
| Intercept | 12.47 | 1 | 0.2459 | 11.94 | 13.01 |  |
| A-*Bacopa monnieri* | 0.0732 | 1 | 0.1922 | -0.3456 | 0.4920 | 1.0000 |
| B-*Acalypha indica* | 0.0882 | 1 | 0.1922 | -0.3306 | 0.5070 | 1.0000 |
| C-*Calotropis Gigantea* | -0.2696 | 1 | 0.1922 | -0.6884 | 0.1492 | 1.0000 |
| AB | 0.1250 | 1 | 0.2511 | -0.4222 | 0.6722 | 1.0000 |
| BC | 0.2500 | 1 | 0.2511 | -0.2972 | 0.7972 | 1.0000 |
| A² | 0.7785 | 1 | 0.1862 | 0.3728 | 1.18 | 1.01 |
| C² | 0.2482 | 1 | 0.1862 | -0.1575 | 0.6539 | 1.01 |

**Note:** The coefficient estimate represents the expected change in response per unit change in factor value when all remaining factors are held constant. The coefficients are adjustments around that average based on the factor settings. When the factors are orthogonal the VIFs are 1; VIFs greater than 1 indicate multi-colinearity, the higher the VIF the more severe the correlation of factors. As a rough rule, VIFs less than 10 are tolerable.
